# Optimizing the balance between heterologous acetate- and CO_2_-reduction pathways in anaerobic cultures of *Saccharomyces cerevisiae* strains engineered for low glycerol production

**DOI:** 10.1101/2023.05.21.541164

**Authors:** Aafke C.A. van Aalst, Ellen H. Geraats, Mickel L.A. Jansen, Robert Mans, Jack T. Pronk

## Abstract

In anaerobic *Saccharomyces cerevisiae* cultures, NADH-cofactor balancing by glycerol formation constrains ethanol yields. Introduction of an acetate-to-ethanol reduction pathway based on heterologous acetylating acetaldehyde dehydrogenase (A-ALD) can replace glycerol formation as ‘redox-sink’ and improve ethanol yields in acetate-containing media. Acetate concentrations in feedstock for first-generation bioethanol production are, however, insufficient to completely replace glycerol formation. An alternative glycerol-reduction strategy bypasses the oxidative reaction in glycolysis by introducing phosphoribulokinase (PRK) and ribulose-1,5-bisphosphate carboxylase (RuBisCO). For optimal performance in industrial settings, yeast strains should ideally first fully convert acetate and, subsequently, continue low-glycerol fermentation via the PRK-RuBisCO pathway. However, anaerobic batch cultures of a strain carrying both pathways showed inferior acetate reduction relative to a strain expressing only the A-ALD pathway. Complete A-ALD-mediated acetate reduction by a dual-pathway strain, grown anaerobically on 50 g L^-1^ glucose and 5 mmol L^-1^ acetate, was achieved upon reducing PRK abundance by a C-terminal extension of its amino-acid sequence. Yields of glycerol and ethanol on glucose were 55% lower and 6% higher, respectively, than those of a non-engineered reference strain. The negative impact of the PRK-RuBisCO pathway on acetate reduction was attributed to sensitivity of the reversible A-ALD reaction to intracellular acetaldehyde concentrations.

## Background

Ethanol, predominantly produced by yeast-based fermentation of renewable carbohydrate feedstocks, can serve as a renewable automotive fuel and as precursor for a range of other products, including ethylene and jet fuel (1, 2). In industrial ethanol production, the sugar feedstock can account for up to 70% of the overall process costs (3). Maximizing the yield of ethanol on sugar is therefore of paramount importance for process economics. The current global fermentative production of ca. 100 Mton ethanol y^-1^ (4) almost exclusively relies on the yeast *Saccharomyces cerevisiae*. Ethanol yields in yeast-based industrial processes can reach up to 92% of the theoretical maximum (5), with loss of substrate carbon primarily due to formation of yeast biomass and glycerol (6).

In anaerobic cultures of wild-type *S. cerevisiae*, glycerol formation via the NAD^+^-dependent glycerol-3-phosphate dehydrogenases Gpd1 and Gpd2 is essential for re-oxidation of ‘surplus’ NADH formed in biosynthetic processes (7, 8). Multiple strategies have been explored to alter the metabolic network of *S. cerevisiae* to reduce NADH formation in biosynthetic reactions and/or rearrange yeast carbon metabolism to couple NADH re-oxidation to the formation of ethanol instead of glycerol (reviewed in (9)). Some of the latter engineering strategies can completely replace glycerol formation in anaerobic laboratory cultures. However, because of the importance of glycerol in yeast osmotolerance (10), application-oriented engineering strategies typically aim at a strong reduction of glycerol formation rather than at its complete elimination (9).

Introduction of a heterologous acetylating acetaldehyde dehydrogenase (A-ALD) into *S. cerevisiae* enables the NADH-dependent reduction of acetyl-CoA to acetaldehyde, which can subsequently be reduced to ethanol by the native yeast alcohol dehydrogenase (11). Coupling this reaction to heterologous pathways involving either phosphoketolase and phosphotransacetylase (12) or pyruvate-formate lyase (13, 14) can couple a net oxidation of NADH to the conversion of glucose to ethanol. Alternatively, in acetate-containing media, A-ALD-expressing yeast strains can use exogenous acetate, which can be activated to acetyl-CoA by native yeast acetyl-CoA synthetase.

Introduction of the *E. coli* A-ALD EutE in *S. cerevisiae*, combined with deletion of *GPD2*, led to a 5-fold lower ratio between glycerol and biomass formation in anaerobic, acetate-supplemented cultures than in the non-engineered parental strain, without affecting specific growth rate (15). However, the combined activity of native acetyl-CoA synthetase (Acs2,(16)), acetaldehyde dehydrogenases (Ald6, Ald5 and Ald4 (17)) and A-ALD can theoretically form an ATP-hydrolysing reaction cycle (18). Additional deletion of *ALD6* is therefore often applied in A-ALD-based redox-cofactor engineering strategies (15, 18, 19).

Development of metabolic engineering strategies aimed at reduction of exogenous acetate to ethanol is predominantly inspired by industrial fermentation of lignocellulosic hydrolysates, in which acetate concentrations can exceed 5 g L^-1^ (20) and negatively affect yeast fermentation (21, 22). In first-generation feedstocks such as corn mash, acetate concentrations of up to 1.2 g L^-1^ are reported (23–25), while glucose concentrations in industrial fermentation processes can reach 300 g L^-1^ (26, 27). During such fermentations, 12-15 g L^-1^ of glycerol is produced (28, 29). To replace the redox-cofactor-balancing role of this amount of glycerol, approximately 4 g L^-1^ of acetate would be required. First-generation feedstocks for ethanol production therefore typically do not contain enough acetate to replace all the glycerol produced.

Ideally, an engineered yeast strain for first-generation processes should reduce all available acetate to ethanol and, after acetate depletion, continue fast anaerobic growth with a low glycerol yield by using another engineered, acetate-independent redox-balancing pathway. An engineered pathway meeting this description is based on bypassing the oxidative reaction in glycolysis by introducing the Calvin-cycle enzymes phosphoribulokinase (PRK) and ribulose-1,5-bisphosphate carboxylase (RuBisCO) (30, 31). Performance of this pathway as a redox-cofactor balancing pathway in yeast was improved by overexpression of structural genes for enzymes of the non-oxidative pentose-phosphate pathway (non-ox PPP↑; p*TDH3*-*RPE1*, p*PGK1*-*TKL1*, p*TEF1*-*TAL1*, p*PGI1*-*NQM1*, p*TPI1*-*RKI1* and p*PYK1*-*TKL2*) and deletion of *GPD2* (31). In addition, multiple copies of an expression cassette for bacterial *cbbM* RuBisCO were integrated in the yeast genome to improve conversion of ribulose-5-bisphosphate into 3-phosphoglycerate (31). While, initially, up to 15 copies of the *cbbM* cassette were integrated (31), a later study indicated that 2 copies were sufficient (32). Expression of a spinach PRK gene from the anaerobically inducible *DAN1* promoter limited toxic effects of PRK during aerobic pre-cultivation (31). The resulting PRK-RuBisCO-synthesizing *S. cerevisiae* strain (IMX2736; Δ*gpd2*, non-ox PPP↑, p*DAN1*-*prk*, 2x *cbbm*, *GroES/GroEL*) showed essentially the same maximum growth rate on glucose as a non-engineered reference strain in anaerobic batch bioreactors while exhibiting a 96% lower glycerol yield and a 10% higher ethanol yield on glucose than the reference strain (32).

The A-ALD and PRK-RuBisCO-based strategies have both been shown to re-oxidize surplus NADH in anaerobic yeast cultures (15, 31) and, in particular in Δ*gpd2* genetic backgrounds, to efficiently compete for NADH with the native glycerol pathway. However, interaction of these two strategies upon their implementation of a single yeast strain has not yet been investigated. The goal of the present study is therefore to study growth and product formation in acetate-containing media of dual-pathway *S. cerevisiae* strains that express both the A-ALD pathway and the PRK-RuBisCO pathway. To this end, engineered strains were grown in anaerobic bioreactor cultures on glucose, using media in which acetate concentrations were either sufficient or insufficient to complete re-oxidation of surplus NADH by acetate reduction. Based on observed patterns of growth and (by)product formation, further engineering was aimed at improving acetate reduction via A-ALD in PRK-RuBisCO-expressing strains.

## Results

### Suboptimal acetate conversion by a yeast strain harbouring both an A-ALD-dependent acetate-reduction and PRK-RuBisCO pathway

*S. cerevisiae Δgpd2* strains expressing a bacterial A-ALD gene can use exogenous acetate as electron acceptor for anaerobic re-oxidation of ‘surplus’ NADH generated in biosynthesis (7). Consistent with earlier reports (15), *S. cerevisiae* IMX2503 (*Δald6 Δgpd2 eutE*) produced 76% less glycerol per amount of biomass formed than the reference strain IME324 (*ALD6 GPD2*) when grown in anaerobic bioreactor batch cultures on 20 g L^-1^ of glucose and 50 mmol L^-1^ acetate (Table 1, Fig. 1A and 1B). This acetate concentration was approximately 5-fold higher than what was calculated to be required for complete re-oxidation of surplus NADH via A-ALD-mediated acetate reduction (9, 11). Specific growth rates of the two strains were not significantly different, but strain IMX2503 showed a 6.7% higher ethanol yield on glucose than the reference strain. Slow, EutE-independent consumption of acetate by the reference strain (Table 1) was previously attributed to the use of acetate-derived acetyl-CoA as a biosynthetic precursor (33).

**Figure 1:**
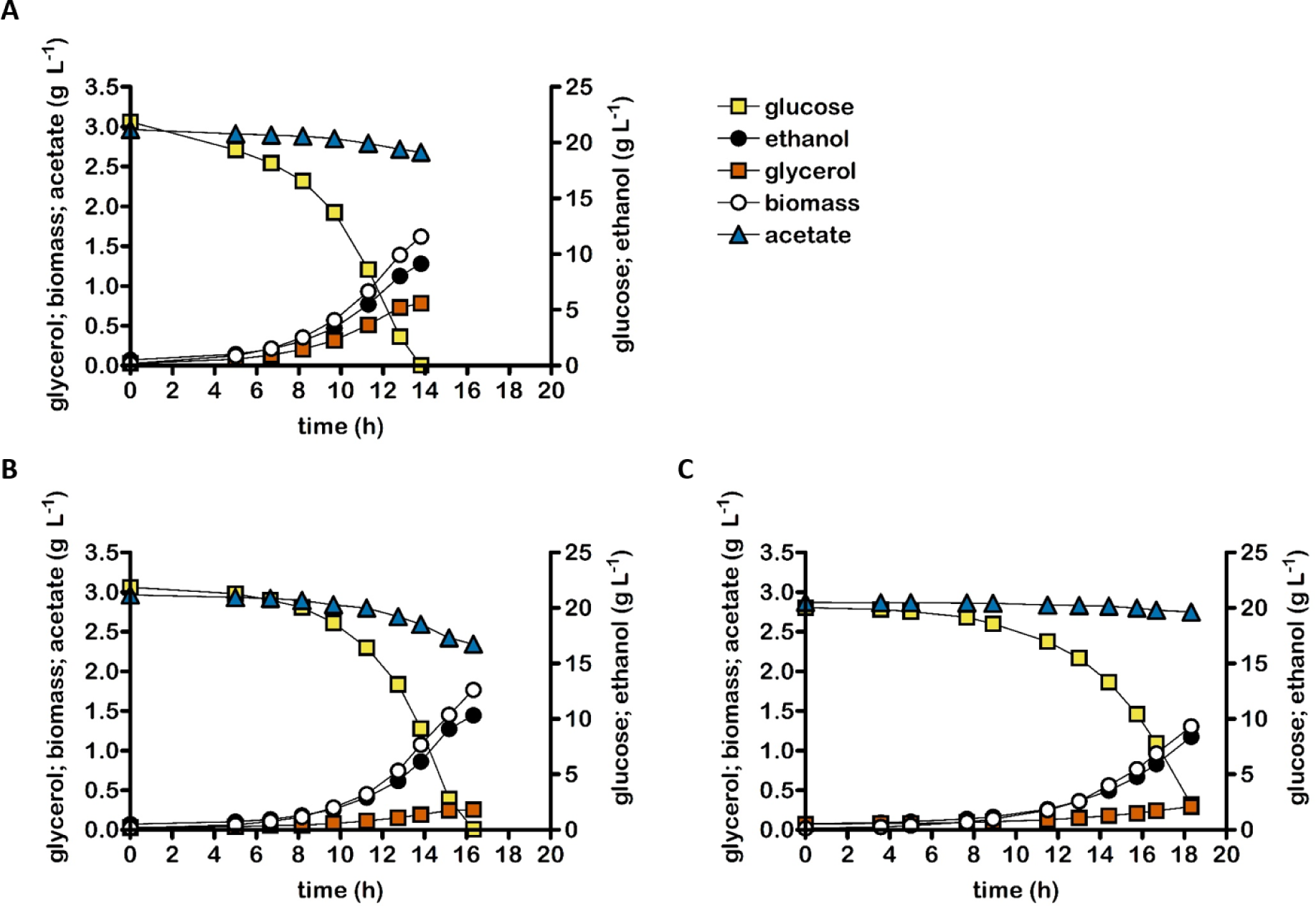
Concentrations of biomass, ethanol, glycerol and acetate in anaerobic bioreactor batch cultures of *S. cerevisiae* strains IME324 (reference strain, **A**), IMX2503 (Δ*gpd2* Δ*ald6* p*TDH3*-*eutE*, **B**), IMX2502 (non-ox PPP↑ Δ*gpd2* Δ*ald6* p*TDH3*-*eutE* p*DAN1*-*prk* 2x p*TDH3*-*cbbm* p*TPI1*-*groES* p*TEF1*-*groEL*, **C**). Cultures were grown anaerobically at pH 5.0 and at 30 °C on synthetic medium containing 20 g L^-1^ glucose and 50 mmol L^-1^ acetate. Non-ox PPP↑ indicates the integration of the expression cassettes of p*TDH3-RPE1*, p*PGK1-TKL1*, p*TEF1-TAL1*, p*PGI1-NQM1*, p*TPI1-RKI1* and p*PYK1-TKL2*. Representative cultures of independent duplicate experiments are shown, corresponding replicate of each culture shown in Additional file 3: Fig. S2.

**Table 1:**
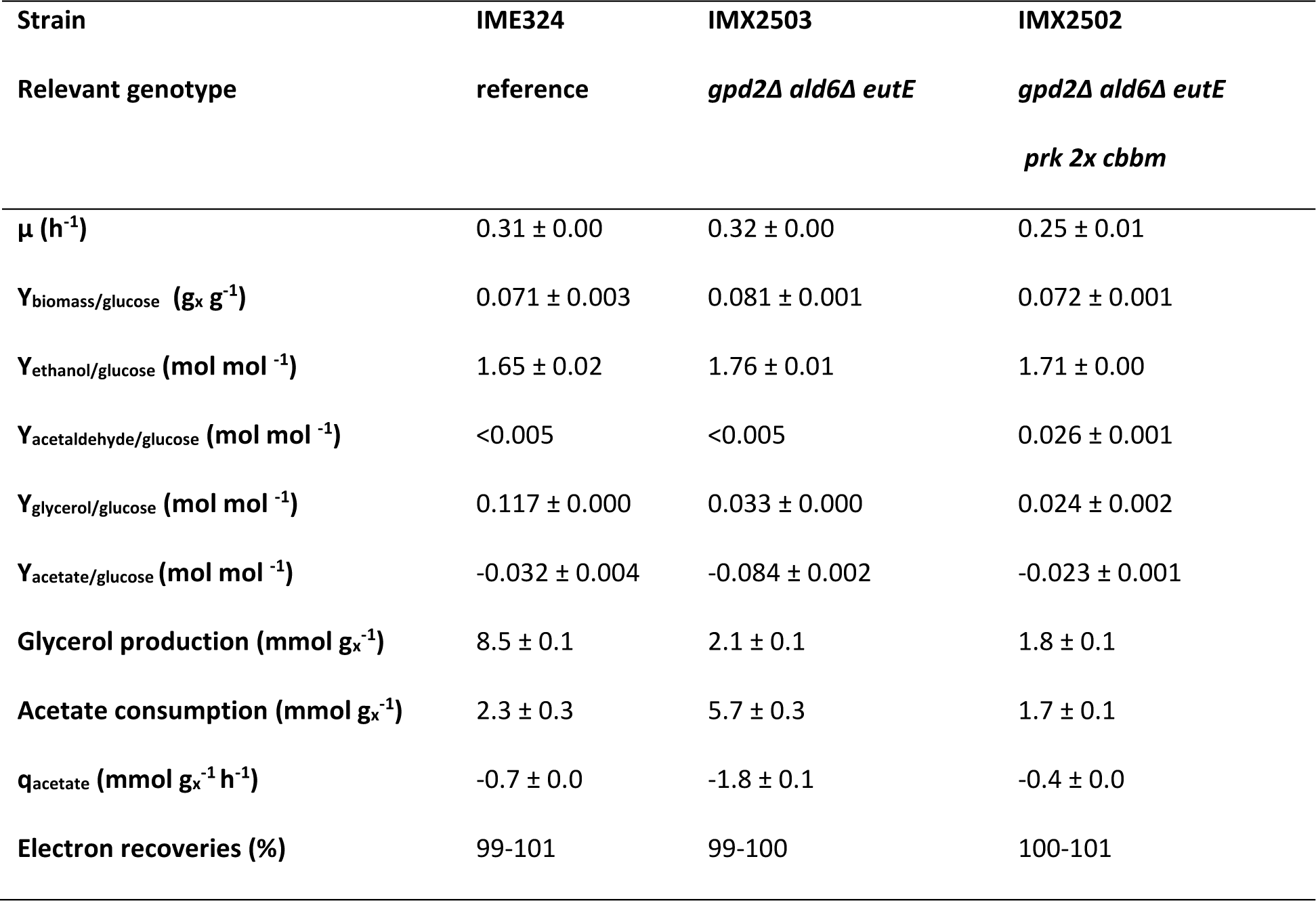
Specific growth rate, stochiometries of biomass and (by)product formation and biomass-specific acetate consumption rates in anaerobic bioreactor batch cultures of *S. cerevisiae* strains IME324 (reference strain), IMX2503 (*Δgpd2 Δald6* p*TDH3*-*eutE*) and IMX2502 (non-ox PPP↑ *Δgpd2 Δald6* p*TDH3*-*eutE* p*DAN1-prk* 2x p*TDH3*-*cbbm* p*TPI1*-*groES pTEF1*-*groEL*). Cultures were grown on synthetic medium with 20 g L^-1^ of glucose and 50 mmol L^-1^ of acetate at pH 5 and at 30 °C and sparged with a 90:10 mixture of N_2_ and CO_2_. Stoichiometries of biomass and metabolite production formation were calculated from at least 6 sampling points in the exponential growth phase, while biomass yield was determined based on biomass and glucose determinations at the first and last time point. Values represent averages ± mean deviations of measurements on independent duplicate cultures. Since the high concentration of CO_2_ in the inlet gas precluded construction of carbon balances, degree-of-reduction balances were used to verify data consistency (34). Non-ox PPP↑ indicates the integration of the expression cassettes of p*TDH3*-*RPE1*, p*PGK1*-*TKL1*, p*TEF1*-*TAL1*, p*PGI1*-*NQM1*, p*TPI1*-*RKI1* and p*PYK1*-*TKL2*. Symbols: μ, specific growth rate; g_x_, gram biomass; Y, yield; q_acetate_: biomass-specific rate of acetate consumption in exponential growth phase.

*S. cerevisiae* IMX2502 (*Δgpd2 Δald6 eutE* non-ox PPP↑ *prk* 2x *cbbm groES/groEL*) combined the genetic modifications in the acetate-reducing strain IMX2503 with introduction of a non-oxidative bypass of glycolysis via phosphoribulokinase (PRK) and ribulose-1,5-bisphosphate carboxylase (RuBisCO, CbbM) (32). Under the conditions described above, this strain showed an even lower rate of acetate consumption than the reference strain IME324 (Table 1). Apparently, simultaneous presence of the two engineered pathways prevented efficient acetate reduction via EutE. However, strain IMX2502 produced 79% less glycerol per amount of biomass than the reference strain IME324, indicating that the PRK-RuBisCO bypass actively contributed to re-oxidation of surplus NADH. Despite its low glycerol formation, strain IMX2502 displayed a mere 4% higher ethanol yield than the reference strain IME324 and a slightly lower ethanol yield than the acetate-reducing strain IMX2503 (Table 1). This increase is approximately 55% lower than theoretically predicted (9) and reported (32) for anaerobic glucose-grown cultures of a congenic ‘PRK-RuBisCO-only’ strain. This lower-than-anticipated ethanol yield of strain IMX2502 coincided with production of up to 2.7 ± 0.1 mmol L^-1^ acetaldehyde, a by-product that was not detected in cultures of the acetate-reducing strain IMX2503 and the reference strain IME324 (Table 1). Formation of acetaldehyde and acetate by slow-growing cultures of PRK-RuBisCO-based *S. cerevisiae* strains was previously attributed to an *in vivo* overcapacity of the non-oxidative bypass of glycolysis (32). The acetaldehyde yield on glucose of strain IMX2502 was 3.6-fold higher than previously reported for anaerobic glucose-grown batch cultures of a congenic PRK-RuBisCO-strain (0.007 ± 0.001 mol acetaldehyde (mol glucose)^-1^, (32)).

### Performance of engineered strains at low acetate-to-glucose ratios

To explore performance of engineered strains at acetate-to-glucose ratios that are more representative for those in first-generation feedstocks such as corn mash, anaerobic bioreactor batch cultures were grown on 50 g L^-1^ of glucose and 5 mmol L^-1^ acetate (Fig. 2, Table 2). Biomass and ethanol yields on glucose of the reference strain IME324 in these cultures were 32% higher and 10% lower, respectively, than in cultures grown at the higher acetate-to-glucose ratio (Tables 1 and 2). These results are in line with a higher maintenance-energy requirement in cultures grown at 50 mmol L^-1^ acetate, caused by weak-acid uncoupling of the plasma-membrane pH gradient (35). Expressed per amount of formed biomass, acetate consumption by strain IME324 was ca. 10-fold lower than in cultures grown at the higher acetate-to-glucose ratio (Tables 1 and 2, respectively) and over two-thirds of the added 5 mmol L^-1^ acetate remained unused (Fig. 2). In contrast, cultures of strain IMX2503 (*Δald6 Δgpd2 eutE*) completely consumed acetate under these conditions (Fig. 2). However, strain IMX2503 showed a mere 12% lower overall glycerol production per amount of biomass formed than the reference strain IME324 (Table 2), as compared to a 76% lower value in cultures grown at a high acetate-to-glucose ratio (Table 1). Consistent with the small difference in glycerol yield, ethanol yields on glucose of the two strains measured in cultures grown at the low acetate-to-glucose ratio were not significantly different. These results are consistent with the dynamics of acetate consumption in the low-acetate cultures of strain IMX2503, which showed a progressive decrease of the biomass-specific rate of acetate consumption during the first phase of batch cultivation (Additional file 3, supplementary Fig. S1). This may reflect the high K_m_ (ca. 8.8 mM) of Acs2, the sole acetyl-coenzyme A synthetase isozyme expressed in anaerobic batch cultures of *S. cerevisiae* (16). The decreasing biomass-specific rate of acetate conversion of strain IMX2503 forced a progressively larger fraction of surplus NADH to be reoxidized via Gpd1-dependent glycerol formation. When, after only approximately half of the glucose had been consumed, acetate was depleted (Fig. 2), this requirement became absolute and, as in wild-type *S. cerevisiae* (29), further anaerobic growth became strictly coupled to glycerol production.

**Figure 2:**
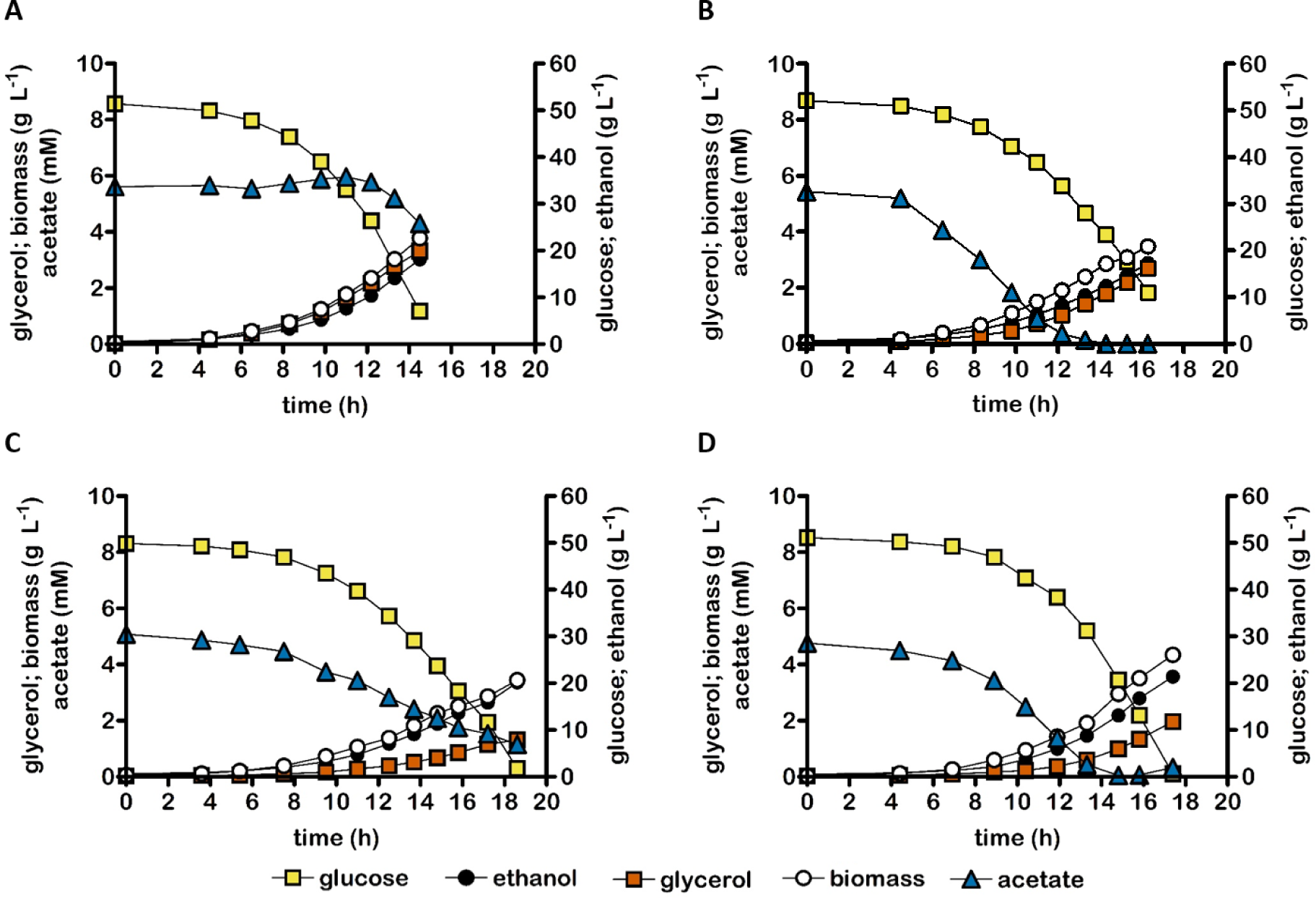
Concentrations of biomass, ethanol, glycerol and acetate in anaerobic bioreactor batch cultures of *S. cerevisiae* strains IME324 (reference strain, **A**), IMX2503 (Δ*gpd2* Δ*ald6 eutE*, **B**), IMX2502 (non-ox PPP↑ Δ*gpd2* Δ*ald6* p*TDH3*-*eutE* p*DAN1*-*prk* 2x p*TDH3*-*cbbm* p*TPI1*-*groES* p*TEF1*-*groEL*, **C**), and IMX2723 (non-ox PPP↑ Δ*gpd2* Δ*ald6* p*TDH3*-*eutE* p*DAN1*-*prk*-19aa (32) 2x p*TDH3*-*cbbm* p*TPI1*-*groES* p*TEF1*-*groEL*, **D**). Cultures were grown anaerobically at pH 5.0 and at 30 °C on synthetic medium containing 50 g L^-1^ glucose and 5.0 mmol L^-1^ acetate. Non-ox PPP↑ indicates the integration of the expression cassettes of p*TDH3*-*RPE1*, p*PGK1*-*TKL1*, p*TEF1*-*TAL1*, p*PGI1*-*NQM1*, p*TPI1*-*RKI1* and p*PYK1*-*TKL2*. Representative cultures of independent duplicate experiments are shown, corresponding replicate of each culture shown in Additional file 3: Fig. S3.

**Table 2:** Stoichiometries of biomass and (by)product formation and biomass, ethanol, glycerol and acetate yield on glucose of anaerobic bioreactor batch cultures of *S. cerevisiae* strains IME324 (reference strain), IMX2503 (*Δgpd2 Δald6* p*TDH3*-*eutE*), IMX2502 (non-ox PPP↑ *Δgpd2 Δald6* p*TDH3*-*eutE* p*DAN1-prk* 2x p*TDH3*-*cbbm* p*TPI1*-*groES pTEF1*-*groEL*) and IMX2723 (non-ox PPP↑ *Δgpd2 Δald6* p*TDH3*-*eutE* p*DAN1-prk*-19aa 2x p*TDH3*-*cbbm* p*TPI1*-*groES pTEF1*-*groEL*). Cultures were grown on synthetic medium with 50 g L^-1^ of glucose and 5.0 mmol L^-1^ acetate at pH 5 and at 30 °C and sparged with a 90:10 mixture of N_2_ and CO_2_. Stoichiometries of biomass and metabolite production were calculated using the first two and last two sampling points, and the stoichiometry of biomass and acetate consumption was calculated using sample points before acetate depletion. Since the high concentration of CO_2_ in the inlet gas precluded construction of accurate carbon balances, degree-of-reduction balances were used to verify data consistency (34). Non-ox PPP↑ indicates the integration of the expression cassettes of p*TDH3*-*RPE1*, p*PGK1*-*TKL1*, p*TEF1*-*TAL1*, p*PGI1*-*NQM1*, p*TPI1*-*RKI1* and p*PYK1*-*TKL2*. Symbols: x, biomass; Y, yield. Values represent averages ± mean deviations of measurements on independent duplicate cultures.

| Strain | IME324 | IMX2503 | IMX2502 | IMX2723 |
| --- | --- | --- | --- | --- |
| Relevant genotype | reference | $\Delta gpd2 \Delta ald6$<br>*eutE* | $\Delta gpd2 \Delta ald6$<br>*eutE prk*<br>*2x cbbm* | $\Delta gpd2 \Delta ald6$<br>*eutE prk-19aa*<br>*(32) 2x cbbm* |
| Y<sub>biomass/glucose</sub> (g<sub>x</sub> g<sup>-1</sup>) | 0.084 ± 0.000 | 0.085 ± 0.000 | 0.073 ± 0.001 | 0.087 ± 0.001 |
| Y<sub>ethanol/glucose</sub> (mol mol<sup>-1</sup>) | 1.55 ± 0.01 | 1.58 ± 0.00 | 1.59 ± 0.02 | 1.64 ± 0.02 |
| Y<sub>glycerol/glucose</sub> (mol mol<sup>-1</sup>) | 0.150 ± 0.000 | 0.125 ± 0.000 | 0.053 ± 0.000 | 0.070 ± 0.002 |
| Y<sub>acetate/glucose</sub> (mol mol<sup>-1</sup>) | -0.004 ± 0.000 | -0.067 ± 0.002 | -0.021 ± 0.002 | -0.053 ± 0.009 |
| Glycerol production<br>(mmol g<sub>x</sub><sup>-1</sup>) | 9.9 ± 0.0 | 8.2 ± 0.0 | 4.3 ± 0.1 | 4.5 ± 0.1 |
| Acetate consumption<br>(mmol g<sub>x</sub><sup>-1</sup>) | 0.2 ± 0.0 | 3.5 ± 0.2 | 1.4 ± 0.2 | 2.5 ± 0.0 |
| Electron recovery | 99-99 | 99-99 | 94-94 | 99-99 |

In cultures of the dual-pathway-strain IMX2502 (*Δgpd2 Δald6 eutE* non-ox PPP↑ *prk* 2x *cbbm groES/groEL*) grown at a low acetate-to-glucose ratio, glycerol production per amount of biomass was 65% and 49% lower, respectively, than in corresponding cultures of the reference strain IME324 and the acetate-reducing strain IMX2503 (Table 2). However, acetate consumption expressed per amount of biomass formed by strain IMX2502 was 2.6-fold lower than in cultures of strain IMX2503 (Table 2) and, upon glucose depletion, not all acetate had been consumed (Fig. 2). A 6 to 7% gap in the degree-of-reduction balance for low-acetate cultures of the dual-pathway-strain IMX2502 (Table 2) probably suggested that, as observed for cultures of this strain grown at a high acetate-to-glucose ratio (Table 1), acetaldehyde was formed as a byproduct. This interpretation was also consistent with an only slightly (2%) higher ethanol yield on glucose and a 14% lower biomass yield relative than observed for the reference strain IME324 (Table 2).

### Tuning the PRK-RuBisCO pathway for improved acetate conversion in an A-ALD-based acetate-reducing yeast strain

The results presented above show that, at both high and low acetate-to-glucose ratios, presence of a functional PRK-RuBisCO pathway in an A-ALD-based acetate-reducing *S. cerevisiae* strain impeded *in vivo* acetate reduction. We recently reported that a 19 amino-acid C-terminal extension (‘19aa’) of the heterologous PRK protein reduced its abundance in engineered yeast strains. This modification was shown to mitigate an overcapacity of the PRK-RuBisCO pathway that led to formation of acetaldehyde and acetate in slow-growing cultures (32). To investigate whether this modification could also affect interference of the PRK-RuBisCO pathway with the A-ALD-dependent reduction of exogenous acetate, strain IMX2723 (*Δgpd2 Δald6 eutE* non-ox PPP↑ p*DAN1-prk*-19aa *cbbm groES/groEL*) was grown in anaerobic batch cultures on 50 g L^-1^ glucose and 50 mmol L^-1^ acetate. In contrast to strain IMX2502, in which the PRK protein did not carry the C-terminal extension, strain IMX2723 consumed all acetate added to the medium and did not show a gap in degree-of-reduction balances or reduced biomass yield (Table 2). In addition, it showed a lower glycerol yield than the acetate-reducing strain IMX2503 and, relative to the reference strain IME324, it showed a 6% higher ethanol yield on glucose.

## Discussion

Previous studies demonstrated that, in *Δgpd2* genetic backgrounds, the A-ALD-based acetate-reduction pathway and the PRK-RuBisCO pathway can each efficiently compete for NADH with the remaining glycerol production pathway and, thereby, enable improved ethanol yields in anaerobic *S. cerevisiae* cultures (15, 31). Here, introduction of both pathways in a single *S. cerevisiae* strain was shown to strongly impede *in vivo* activity of acetate reduction via the A-ALD pathway. In addition, anaerobic batch cultures of the dual-pathway strain produced high levels of acetaldehyde, a byproduct that was previously found in slow-growing cultures of strains carrying the PRK-RuBisCO pathway (32).

Acetaldehyde produced by the PRK-RuBisCO pathway might influence *in vivo* activity of the reversible A-ALD reaction (ΔG^0^’ = 17.6 kJ mol^-1^ for the reductive reaction (36). Its reactants NADH/NAD^+^ and acetyl-CoA/CoA are conserved moieties, whose concentration ratios are likely to be constrained by their involvement in a large number of metabolic reactions. An estimate of the maximum permissive acetaldehyde concentration for acetyl-CoA reduction (ΔG_R_^’^ = 0), based on reported NADH:NAD^+^ and acetyl-CoA:CoA ratios in glucose-grown yeast cultures (Additional file 3), yielded a value of 0.35 mmol L^-1^. Assuming that acetaldehyde diffuses freely out of the yeast cell, the acetaldehyde concentration measured in the fermentation broth was assumed to be the same as the intracellular acetaldehyde concentration. Acetaldehyde concentrations in glucose-acetate grown anaerobic batch cultures of the dual pathway strain IMX2502 were an order of magnitude higher (1.0 to 2.8 mmol L^-1^, Additional file 1) than this estimated value. In contrast, acetaldehyde concentrations in cultures of the ‘single-pathway’ acetate-reducing strain IMX2503 did not exceed 0.35 mmol L^-1^ (Additional file 1). This analysis supports the interpretation that the elevated acetaldehyde concentrations in cultures of the dual-pathway strain rendered a net reduction of acetyl-CoA to acetaldehyde by A-ALD thermodynamically impossible.

Acetaldehyde concentrations in anaerobic batch cultures of the dual-pathway strain IMX2502 exceeded those reported for anaerobic batch cultures of congenic ‘PRK-RuBisCO-only’ strains (32). This difference may be related to the deletion of *ALD6*, which encodes cytosolic NADP^+^-dependent acetaldehyde dehydrogenase. *ALD6* was deleted in A-ALD-containing strains, including the dual pathway strain, to prevent a cytosolic ATP-dissipating futile cycle, consisting of A-ALD, Ald6 and the acetyl-CoA synthetase. This futile cycle was implicated in delayed growth of A-ALD-based strains in high-osmolarity media (18). In the dual-pathway context, eliminating a key enzyme for acetaldehyde conversion may be less desirable and further research is needed to investigate whether and how expression levels of *ALD6* affect acetaldehyde production by PRK-RuBisCO-based strains.

To simulate the low acetate-to-glucose ratios in first-generation feedstocks for industrial ethanol production, we used a medium containing 50 g L^-1^ glucose and 5 mmol L^-1^ acetate (corresponding to 0.3 g L^-1^ acetic acid). At this low initial concentration of acetate, the biomass-specific rate of acetate conversion by strain IMX2503 (*Δgpd2 Δald6 eutE*) declined as the acetate concentration decreased (Additional file 3, supplementary Fig. S1), which indicated a suboptimal affinity of this strain for acetate. While acetate concentrations in industrial media can be 3- to 4-fold higher (23–25), improvement of the kinetics of acetate reduction may be required for fast and complete acetate conversion. This could for example be achieved via expression of acetyl-CoA synthetases with better affinity for acetate than the anaerobically expressed isoenzyme Acs2. A candidate protein could be the native Acs1, which is not synthesized under anaerobic conditions and has a 30-fold lower Km than Acs2 (0.32 and 8.8 mmol L^-1^, respectively (16)). However, Acs1 is subject to glucose catabolite inactivation (37), which complicates its application in glucose-grown batch cultures. Alternatively, highly active heterologous acetyl-CoA synthetases, such as an optimized variant of the *Salmonella enterica* enzyme (38–40) may be applied. Alternatively, strains with improved affinity for acetate, potentially also due to changes in acetate transport across the plasma membrane, may be obtained by laboratory evolution (41). Such experiments could for example be based on anaerobic, acetate-limited chemostat cultures of A-ALD-expressing *Δgpd1 Δgpd2* strains (42).

A C-terminal extension of the heterologous PRK was previously shown to reduce acetaldehyde production in slow-growing cultures of PRK-RuBisCO-carrying *S. cerevisiae* (32). The same strategy for ‘tuning’ activity of the PRK-RuBisCO pathway enabled complete conversion of acetate in media with low acetate-to-glucose ratios, while maintaining a low glycerol yield and high ethanol yield after acetate depletion. This result provides a first proof-of-principle for efficient conversion of feedstocks for industrial bioethanol production with a low acetate content, thereby preventing continual increase of acetate via recycle water (43) and improving ethanol yield. However, controlling pathway activity by modification of the abundance of a single enzyme is likely to be a too static approach for application under industrial conditions. Strategies to dynamically regulate expression of the PRK-RuBisCO and A-ALD pathways in response to changes in medium composition could, for example, be based on the design and construction of synthetic regulatory loops based on prokaryotic sensor proteins for acetaldehyde and acetate, such as *Bacillus subtilis* AlsR, which activates transcription in response to acetate (44, 45).

## Materials and Methods

### Growth media and strain maintenance

*S. cerevisiae* strains constructed and/or used in this study (Table 4) all originate from the CEN.PK lineage (46, 47). Yeast strains were propagated in YPD medium (10 g L^-1^ Bacto yeast extract (Thermo Fisher Scientific, Waltham MA), 20 g L^-1^ Bacto^TM^ peptone (Thermo Fisher Scientific), 20 g L^-1^ glucose) or synthetic medium with vitamins (SM; 3.0 g L^-1^ KH_2_PO_4_, 0.5 g L^-1^ MgSO_4_·7H_2_O, 5.0 g L^-1^ (NH_4_)_2_SO_4_, (48)) prepared and sterilized as described previously. Concentrated solutions of glucose were autoclaved separately for 20 min at 110 °C and supplemented to SM to a final concentration of 20 g L^-1^ or 50 g L^-1^. For growth of uracil-auxotrophic strains, 150 mg L^-1^ uracil was added to SM by adding a concentrated uracil solution (3.75 g L^-1^) autoclaved at 120 °C for 20 min (49). To select for presence of an acetamidase marker cassette (50), (NH_4_)_2_SO_4_ was replaced by 6.6 g L^-1^ K_2_SO_4_ and 0.6 g L^-1^ filter-sterilized acetamide. Where indicated, acetic acid (≥99.8%, Honeywell, Charlotte NC) was added to media to a concentration of 0.3 g L^-1^ or 3 g L^-1^. A concentrated stock solution of Tween 80 (polyethylene glycol sorbitan monoelate; Merck, Darmstadt, Germany) and ergosterol (98%; Acros Organics-Thermo Fisher Scientific) (420 g L^-1^ Tween and 10 g L^-1^ ergosterol) was prepared in absolute ethanol (Supelco: Sigma-Aldrich, St. Louis MI). For anaerobic cultivation, 1 mL of this solution was added per litre of medium (51). *Escherichia coli* XL1-Blue stock cultures were propagated in lysogeny broth (LB) medium (52). For strain maintenance, glycerol (30% v/v final concentration) was added to late exponential phase cultures and stored at −80 °C. Solid media were prepared by adding 20 g L^-1^ Bacto agar (Becton Dickinson, Breda, The Netherlands) to mineral salts solutions. Vitamin solution, glucose and when required acetamide were added to SM-agar media after cooling to 60 °C. *S. cerevisiae* cultures on agar plates were incubated at 30 °C until colonies appeared (1-5 days) and *E. coli* cultures on plates were incubated overnight at 37 °C.

### Construction of plasmids and expression cassettes

DNA fragments for construction of plasmids and expression cassettes were PCR amplified with Phusion High Fidelity DNA Polymerase (Thermo Fisher Scientific) according to the manufacturer’s manual. Diagnostic colony PCR was performed using DreamTaq polymerase (Thermo Fisher Scientific). DNA fragments were separated by electrophoresis on 1% (w/v) agarose (Sigma-Aldrich) gels in 1xTAE (40 mM Tris-acetate pH 8.0 and 1 mM EDTA). Fragments were isolated from gels with the Zymoclean Gel DNA Recovery kit (Zymo Research, Irvine CA) or isolated from PCR mixes with a GeneJET kit (Thermo Fisher Scientific). DNA concentrations were measured with a NanoDrop 2000 spectrophotometer (wavelength 260 nm; Thermo Fisher Scientific). Plasmid assembly was performed by *in vitro* Gibson assembly using a HiFi DNA Assembly Master Mix (New England Biolabs, Ipswich MA), downscaled to 5 µL reaction volume. 1 µL of reaction mixture was used for heat-shock transformation (53) of *E. coli* XL-1 Blue. Plasmids were isolated from *E. coli* XL-I Blue transformants with the Sigma GenElute Plasmid Kit (Sigma-Aldrich) according to the manufacturer’s instructions. Plasmids used and constructed in this study are listed in Table 3 and oligonucleotide primers are listed in Table S1.

**Table 3:** Plasmids used in this study. *Kl* denotes *Kluyveromyces lactis*

| Plasmid name | Characteristics | Reference |
| --- | --- | --- |
| **p426_TEF** | 2 µm ori, *URA3*, pTEF1-tCYC1 (empty vector) | (54) |
| **pMEL10** | 2 µm ori, *KIURA*, gRNA-CAN1.Y | (55) |
| **pMEL11** | 2 µm ori, *AmdS*, gRNA-CAN1.Y | (55) |
| **pROS10** | 2 µm ori, *KIURA3*, gRNA-CAN1.Y gRNA-ADE2.Y | (55) |
| **pUDI076** | pRS406-*TDH3*p-*eutE*-CYC1t | (18) |
| **pUDR103** | 2 µm ori, *KIURA3*, gRNA-SGA1.Y | (31) |
| **pUDR264** | 2 µm ori, *AmdS*, gRNA.ALD6.Y | (18) |
| **pUDR744** | 2 µm ori, *KIURA3*, gRNA.GPD2.Y gRNA.GPD2.Y | This work |

A unique Cas9-recognition sequence in *GPD2* was identified as described previously (55). pUDR744 was constructed by PCR amplification of pROS10 with primer 5793 to obtain a linear backbone, and PCR amplification of pROS10 with primer 7839 to obtain the insert fragment containing the *GPD2* gRNA. The plasmid was then assembled by *in vitro* Gibson Assembly. A p*TDH3-eutE-*t*CYC1* integration cassette was obtained by PCR amplification with primers 16615/16616 with pUDI076 as template, adding 60 bp terminal sequences homologous to sequences directly upstream and downstream of the coding region of *ALD6*. A dsDNA repair fragment for *GDP2* deletion was obtained by mixing primers 6969/6970 in a 1:1 molar ratio, heating the mixture to 95 °C for 5 min and subsequently cooling down at room temperature.

#### Yeast genome editing

The lithium-acetate method (56) was used for yeast transformation. Correct Cas9-mediated integration or deletion was routinely verified by diagnostic PCR and single-colony isolates were restreaked thrice on SMD (SM with 20 g L^-1^ glucose) and stored at −80 °C.

**Table 4:**
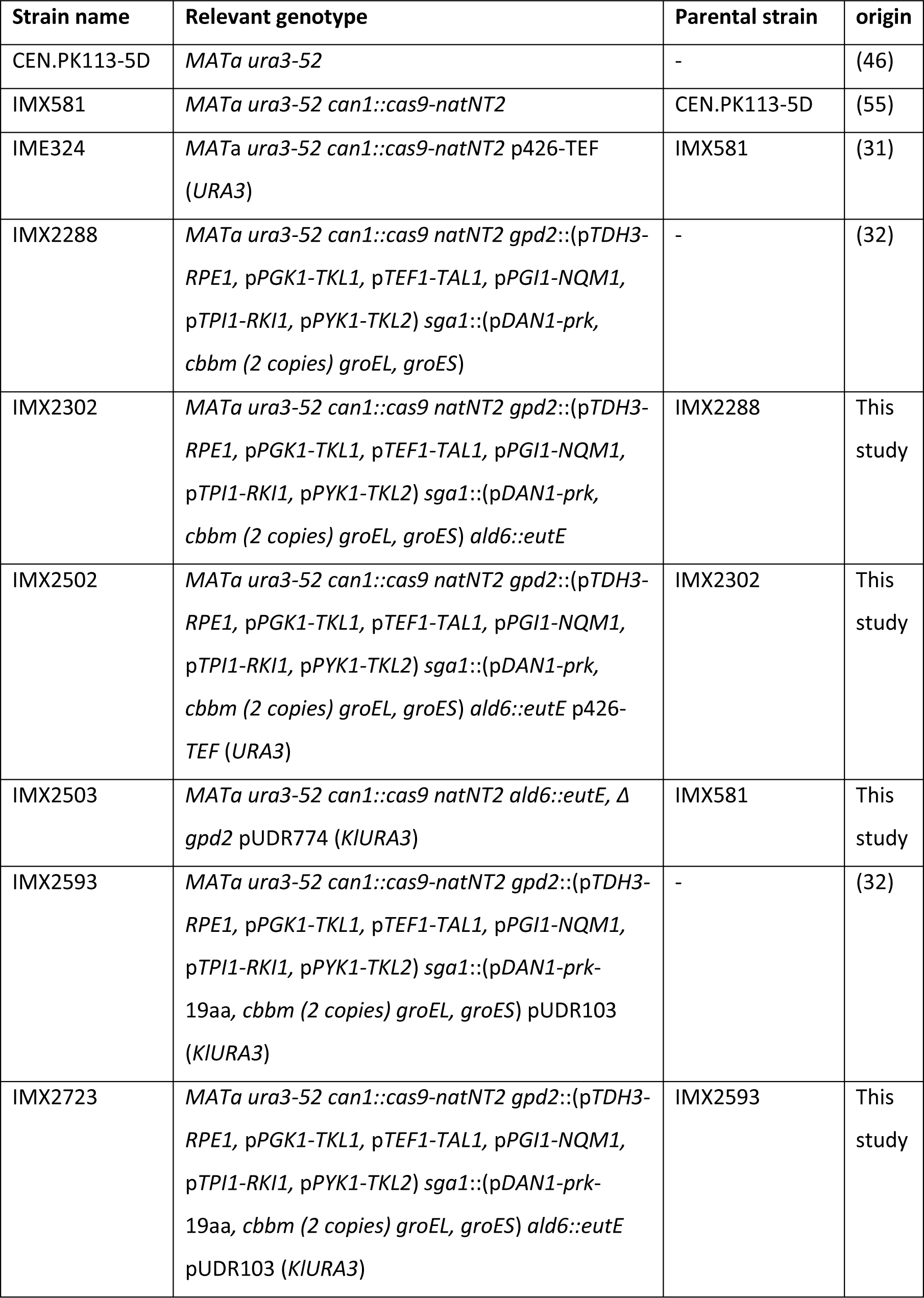
*S. cerevisiae* strains used in this study. *Kl* denotes *Kluyveromyces lactis*

| Strain name | Relevant genotype | Parental strain | origin |
| --- | --- | --- | --- |
| CEN.PK113-5D | *MATa ura3-52* | - | (46) |
| IMX581 | *MATa ura3-52 can1::cas9-natNT2* | CEN.PK113-5D | (55) |
| IME324 | *MATa ura3-52 can1::cas9-natNT2* p426-TEF<br>(*URA3*) | IMX581 | (31) |
| IMX2288 | *MATa ura3-52 can1::cas9 natNT2 gpd2::(pTDH3-RPE1, pPGK1-TKL1, pTEF1-TAL1, pPGI1-NQM1, pTPI1-RKI1, pPYK1-TKL2) sga1::(pDAN1-prk, cbbm (2 copies) groEL, groES)* | - | (32) |
| IMX2302 | *MATa ura3-52 can1::cas9 natNT2 gpd2::(pTDH3-RPE1, pPGK1-TKL1, pTEF1-TAL1, pPGI1-NQM1, pTPI1-RKI1, pPYK1-TKL2) sga1::(pDAN1-prk, cbbm (2 copies) groEL, groES) ald6::eutE* | IMX2288 | This study |
| IMX2502 | *MATa ura3-52 can1::cas9 natNT2 gpd2::(pTDH3-RPE1, pPGK1-TKL1, pTEF1-TAL1, pPGI1-NQM1, pTPI1-RKI1, pPYK1-TKL2) sga1::(pDAN1-prk, cbbm (2 copies) groEL, groES) ald6::eutE* p426-TEF (*URA3*) | IMX2302 | This study |
| IMX2503 | *MATa ura3-52 can1::cas9 natNT2 ald6::eutE, Δgpd2* pUDR774 (*KIURA3*) | IMX581 | This study |
| IMX2593 | *MATa ura3-52 can1::cas9-natNT2 gpd2::(pTDH3-RPE1, pPGK1-TKL1, pTEF1-TAL1, pPGI1-NQM1, pTPI1-RKI1, pPYK1-TKL2) sga1::(pDAN1-prk-19aa, cbbm (2 copies) groEL, groES)* pUDR103 (*KIURA3*) | - | (32) |
| IMX2723 | *MATa ura3-52 can1::cas9-natNT2 gpd2::(pTDH3-RPE1, pPGK1-TKL1, pTEF1-TAL1, pPGI1-NQM1, pTPI1-RKI1, pPYK1-TKL2) sga1::(pDAN1-prk-19aa, cbbm (2 copies) groEL, groES) ald6::eutE* pUDR103 (*KIURA3*) | IMX2593 | This study |

*S. cerevisiae* IMX2302 was constructed by co-transforming strain IMX2288 with pUDR264 and repair fragment p*TDH3-eutE-*t*CYC1* (with homologous flanks to the upstream and downstream sequence of *ALD6* open reading frame). Transformants were selected on SMD with acetamide and uracil. To restore uracil prototrophy, strain IMX2302 was transformed with the *URA3*-carrying plasmid p426-*TEF* (empty vector), yielding strain IMX2502.

Strain IMX2503 was constructed by co-transforming IMX581 with gRNA-plasmids pUDR744 (targeting *GPD2*) and pUDR264 (targeting *ALD6*) and repair fragments for *GPD2* deletion (dsDNA homologous to the upstream and downstream sequence of *GPD2* open reading frame) and p*TDH3-eutE-*t*CYC1* (with homologous flanks to the upstream and downstream sequence of *ALD6* open reading frame). Transformants were selected on SMD with acetamide. pUDR264 was removed by selection on SMD, while pUDR744 was retained to support uracil prototrophy.

Strain IMX2723 was obtained from IMX2593 by co-transforming with gRNA-plasmid pUDR264 and repair fragment p*TDH3-eutE-*t*CYC1*. Transformants were selected on SMD with acetamide. pUDR264 was removed by selection on SMD, while pUDR103 was retained to support uracil prototrophy.

### Aerobic shake-flask cultivation

Shake-flask cultures were grown at 30 °C in 500-mL round-bottom shake-flasks containing 100 mL medium, placed in an Innova incubator shaker (Eppendorf Nederland B.V., Nijmegen, The Netherlands) and shaken at 200 rpm.

### Bioreactor cultivation

Anaerobic bioreactor batch cultivation was conducted at 30 °C in 2-L bioreactors (Applikon Getinge, Delft, The Netherlands), with a working volume of 1 L. Culture pH was kept constant at 5.0 by automatic addition of 2 M KOH. All bioreactor cultures were grown on synthetic medium supplemented with vitamins, glucose, acetic acid, the anaerobic growth factors Tween 80 (420 mg L^-1^) and ergosterol (10 mg L^-1^), and 0.2 g L^-1^ antifoam C (Sigma-Aldrich). Cultures were sparged at 0.5 L min^-1^ with an N_2_/CO_2_ (90/10%) gas mixture. The outlet gas stream was cooled to 4 °C in a condenser to minimize evaporation. Oxygen diffusion was minimized by use of Norprene tubing (Saint-Gobain, Amsterdam, The Netherlands) and Viton O-rings (ERIKS, Haarlem, The Netherlands)(51). Inocula were prepared in 500-mL shake flasks containing 100 mL SMD. A first starter culture was inoculated with a frozen stock culture, grown aerobically for 15-18 h at 30 °C and used to inoculate precultures on SMD. Upon reaching mid-exponential phase (OD_660_ of 3-5), these were used to inoculate bioreactor cultures at an initial OD_660_ of 0.25-0.40.

### Analytical methods

Growth was monitored by biomass dry weight measurements (30) and by measuring optical density at 660 nm (OD_660_) on a Jenway 7200 spectrophotometer. Metabolite concentrations were determined by high-performance liquid chromatography (30). A first-order evaporation rate constant of 0.008 h^-1^ was used to correct ethanol concentrations for evaporation (30). Acetaldehyde concentrations were determined in the off-gas and the broth by derivatization using 2,4-DNPH as described previously (32). As carbon recoveries could not be accurately calculated due to the high concentration of CO_2_ in the inlet gas of bioreactor cultures, electron recoveries based on degree of reduction of relevant compounds (34) were used instead.

## Declarations

### Supplementary information

**Additional file 1_supplementary data.** Measurement data of anaerobic batch cultures of IME324, IMX2503 and IMX2502 on 20 g L^-1^ of glucose and 3 g L^-1^ of acetate. Data was used to prepare Table 1 and Figure 1.

**Additional file 2_supplementary data.** Measurement data of anaerobic batch cultures of IME324, IMX2503, IMX2502 and IMX2723 on 50 g L^-1^ of glucose and 0.3 g L^-1^ of acetate. Data was used to prepare Table 2, Figure 2 and Figure S1.

**Additional file 3_supplementary data. Additional information S1.** Description of the calculation to estimate the acetaldehyde concentration at which A-ALD can no longer operate in the reductive direction. **Table S1.** Oligonucleotides used in this study. **Figure S1**. Biomass specific acetate consumption rate at different acetate concentrations of anaerobic batch cultures of IMX2503 on 50 g L^-1^ of glucose and 0.3 g L^-1^ of acetate. **Figure S2**. Growth, glucose consumption, ethanol, acetate and glycerol formation in anaerobic bioreactor batch cultures of *S. cerevisiae* strains IME324, IMX2503 and IMX2502 on 20 g L^-1^ glucose and 50 mmol L^-1^ acetate. **Figure S3**. Growth, glucose consumption, ethanol, acetate and glycerol formation in anaerobic bioreactor batch cultures of *S. cerevisiae* strains IME324, IMX2503, IMX2502 and IMX2723 on 50 g L^-1^ glucose and 5.0 mmol L^-1^ acetate.

## Data availability

Short read DNA sequencing data of the *Saccharomyces cerevisiae* strain IMX2503 were deposited at NCBI under BioProject accession number PRJNA972872. All measurement data used to prepare Figure 1, Figure 2, Table 1 and Table 2 of the manuscript and Figure S1, Figure S2 and Figure S3 of the supplementary materials are available in Additional file 1 and Additional file 2.

## Funding

This work was supported by DSM Bio-based Products & Services B.V. (Delft, The Netherlands). Royal DSM owns intellectual property rights of technology discussed in this paper.

### Authors’ contributions

**AA:** Validation, Methodology, Formal analysis, Investigation, Writing – Original Draft, Writing – Review & Editing, Visualization. **EG**: Investigation, Writing – Review & Editing. **MJ:** Writing – Review & Editing. **RM:** Conceptualization, Supervision, Writing – Review & Editing. **JP:** Conceptualization, Supervision, Writing – Original Draft, Writing – Review & Editing. All authors read and approved the final manuscript.

