## Additional file 3 for "Optimizing the balance between heterologous acetate- and CO_2_-reduction pathways in anaerobic cultures of *Saccharomyces cerevisiae* strains engineered for low glycerol production"

**Supplementary data**

**Additional information S1**

Here, we estimate the threshold concentration of acetaldehyde at which A-ALD can operate in the reductive direction (equation I). For this estimation, we used equation II. By implementing a zero Gibbs free energy change (i.e. thermodynamic equilibrium, so no net reduction or oxidation), equation II was rewritten to equation III. We used a ΔG^0’^ of 17600 J mol^-1^ (1), R is 8.3145 J mol^-1^ K^-1^ and T is 303.15 K. For the CoA:acetyl-CoA-ratio, a value of 3.8 was used based on measurements from aerobic batch cultures of *S. cerevisiae* (2). A value of 10.1 was used for the NAD^+^:NADH ratio, based on a published estimate for anaerobic *S. cerevisiae* cultures (3). This approach yielded a threshold concentration of 0.35 mM acetaldehyde (i.e., based on the assumed CoA:acetyl-CoA and NAD^+^:NADH ratios, A-ALD cannot operate in yeast cells in the direction indicated in equation 1 when the acetaldehyde concentration exceeds 0.35 mM).

$acetyl-CoA+NADH+H^{+}\to acetaldehyde+CoA+NAD^{+}$	(equation I)

$\Delta_{R,red}G= \Delta_{R,red}G^{0'}+RT\ln\left( [acetaldehyde]*\frac{[NAD^{+}]}{[NADH]}*\frac{[CoA]}{[acetyl-CoA]} \right)$	(equation II)

$\left[ acetaldehyde \right]=e^{-\frac{\Delta_{R,red}G^{0^{'}}}{RT}}*\frac{[NADH]}{[NAD^{+}]}*\frac{[acetyl-CoA]}{[CoA]}$	(equation III)

**Table S1: Oligonucleotide primers used in this study.** HPLC refers to High Pressure Liquid Chromatography; PAGE denotes PolyAcrylamide Gel Electrophoresis; DST denotes desalted.

| Primer | Sequence | purification |
| --- | --- | --- |
| 5792 | GTTTTAGAGCTAGAAATAGCAAGTTAAAATAAG | PAGE |
| 5793 | GATCATTTATCTTTCACTGCGGAG | PAGE |
| 5979 | TATTGACGCCGGGCAAGAGC | HPLC |
| 5980 | CGACCGAGTTGCTCTTG | HPLC |
| 6969 | GTATTTTGGTAGATTCAATTCTCTTTCCCTTTCCTTTTCCTTCGCTCCCCTTCCTTATCAAACCAATTTATCATTATACACAAGTTCTACAACTACTACTAGTAACATTACTACAGTTAT | DST |
| 6970 | ATAACTGTAGTAATGTTACTAGTAGTAGTTGTAGAACTTGTGTATAATGATAAATTGGTTTGATAAGGAAGGGGAGCGAAGGAAAAGGAAAGGGAAAGAGAATTGAATCTACCAAAATAC | DST |
| 7610 | GTTGATAACGGACTAGCCTTATTTTAACTTGCTATTTCTAGCTCTAAAACAATTCAGAGCTGTTAGCCATGATCATTTATCTTTCACTGCGGAGAAGTTTCGAACGCCGAAACATGCGCA | PAGE |
| 7839 | TGCGCATGTTTCGGCGTTCGAAACTTCTCCGCAGTGAAAGATAAATGATCATTATGAACTACATCGATATGTTTTAGAGCTAGAAATAGCAAGTTAAAATAAG | PAGE |
| 16615 | TAGAAGAAAAAACATCAAGAAACATCTTTAACATACACAAACACATACTATCAGAAT  ACACACAGGAAACAGCTATGACC | DST |
| 16616 | CAAGTAAGTTTATATGAAAGTATTTTGTGTATATGACGGAAAGAAATGCAGGTTGGTACAGCCGCAAATTAAAGCCTTCG | DST |

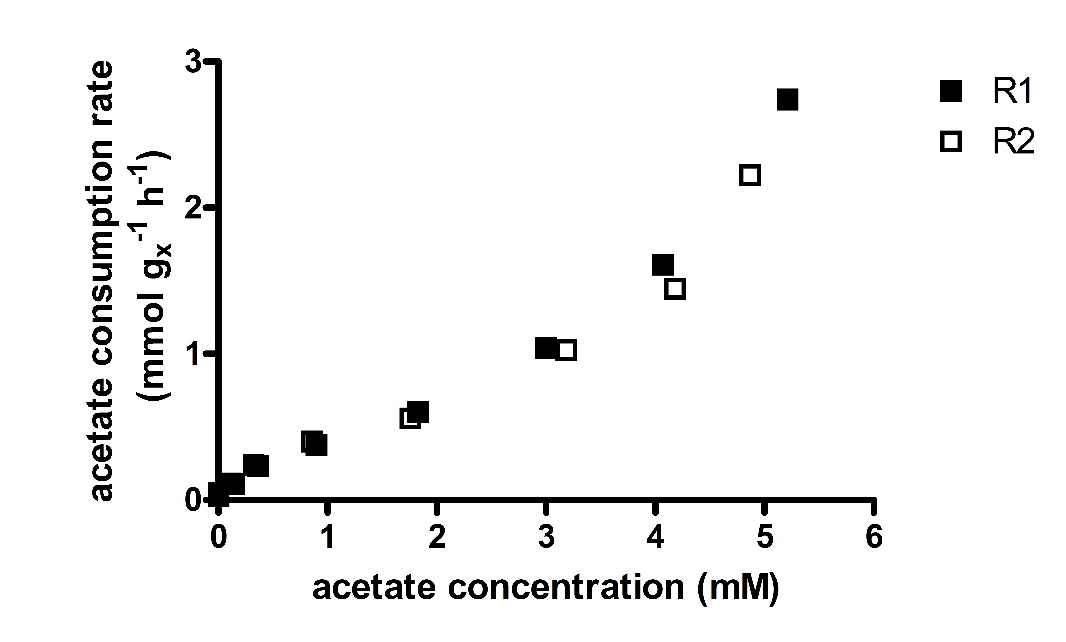

**Figure S1: B**iomass-specific acetate-consumption rate at different acetate concentrations in anaerobic bioreactor batch cultures of S. cerevisiae strain IMX2503 (∆gpd2 ∆ald6 eutE). Cultures were grown on synthetic medium containing 50 g L^-1^ of glucose and 5 mmol L^-1^ of acetate, at pH 5.0, 30°C and sparged with a 90:10 mixture of N_2_ and CO_2_. Cultures were grown in duplicate, represented by R1 and R2. Symbol: g_x_, gram biomass. The biomass-specific acetate-consumption rates were calculated by fitting third-power polynomial spline through a plot of the acetate concentrations at each timepoint. The derivative of the function was used to calculate the acetate-consumption rate at each acetate concentration, which was subsequently normalized using the biomass concentration at each timepoint to obtain the biomass-specific acetate consumption rates.

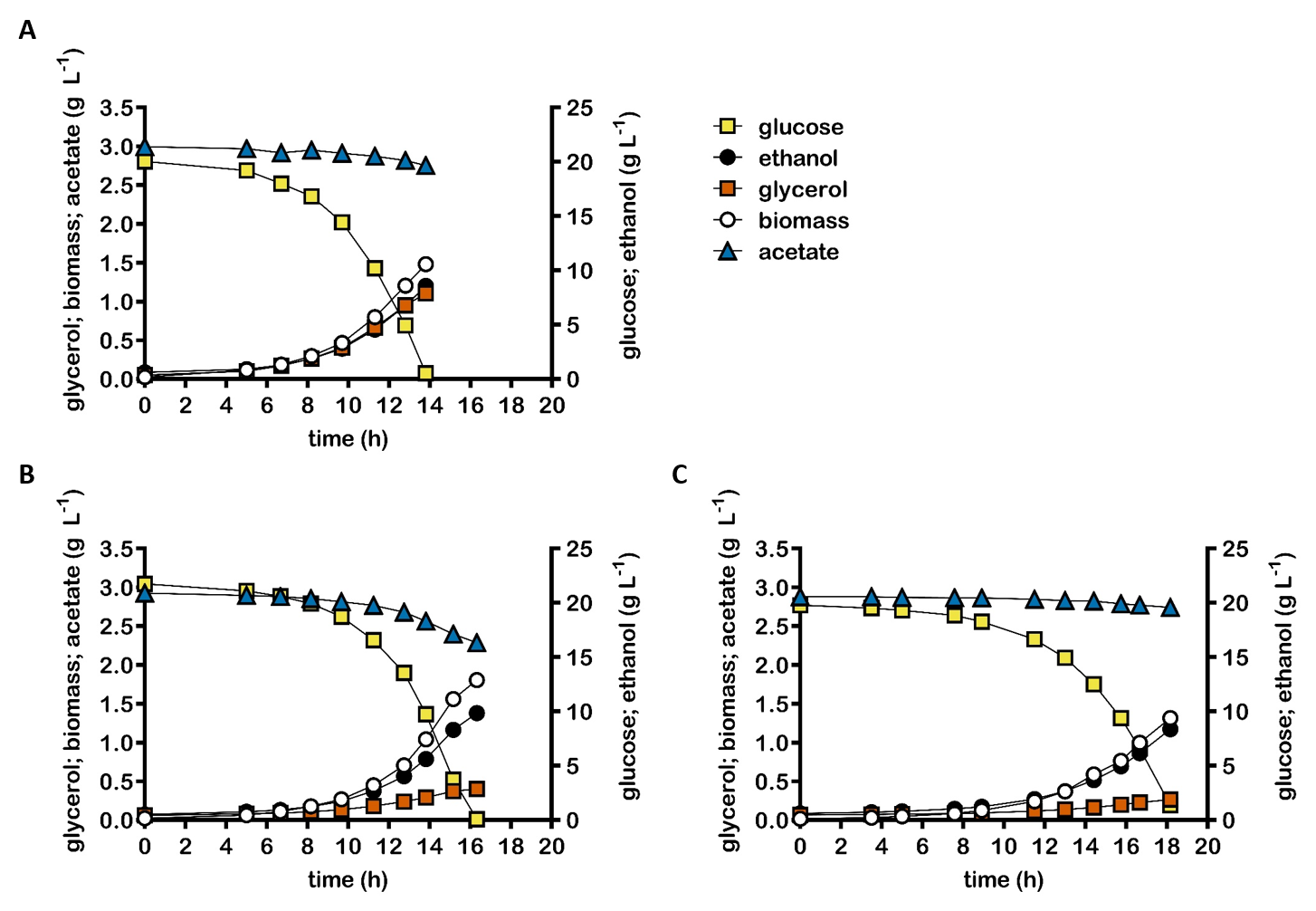

**Figure S2:** Concentrations of biomass, ethanol, glycerol and acetate in anaerobic bioreactor batch cultures of S. cerevisiae strains IME324 (reference strain, **A**), IMX2503 (Δgpd2 Δald6 pTDH3-eutE, **B**), IMX2502 (non-ox PPP↑ Δgpd2 Δald6 pTDH3-eutE pDAN1-prk 2x pTDH3-cbbm pTPI1-groES pTEF1-groEL, **C**). Cultures were grown anaerobically at pH 5.0 and at 30 °C on synthetic medium containing 20 g L^-1^ glucose and 50 mmol L^-1^ acetate. Non-ox PPP↑ indicates the integration of the expression cassettes of pTDH3-RPE1, pPGK1-TKL1, pTEF1-TAL1, pPGI1-NQM1, pTPI1-RKI1 and pPYK1-TKL2. Representative cultures of independent duplicate experiments are shown, corresponding replicate of each culture shown in Figure 1.

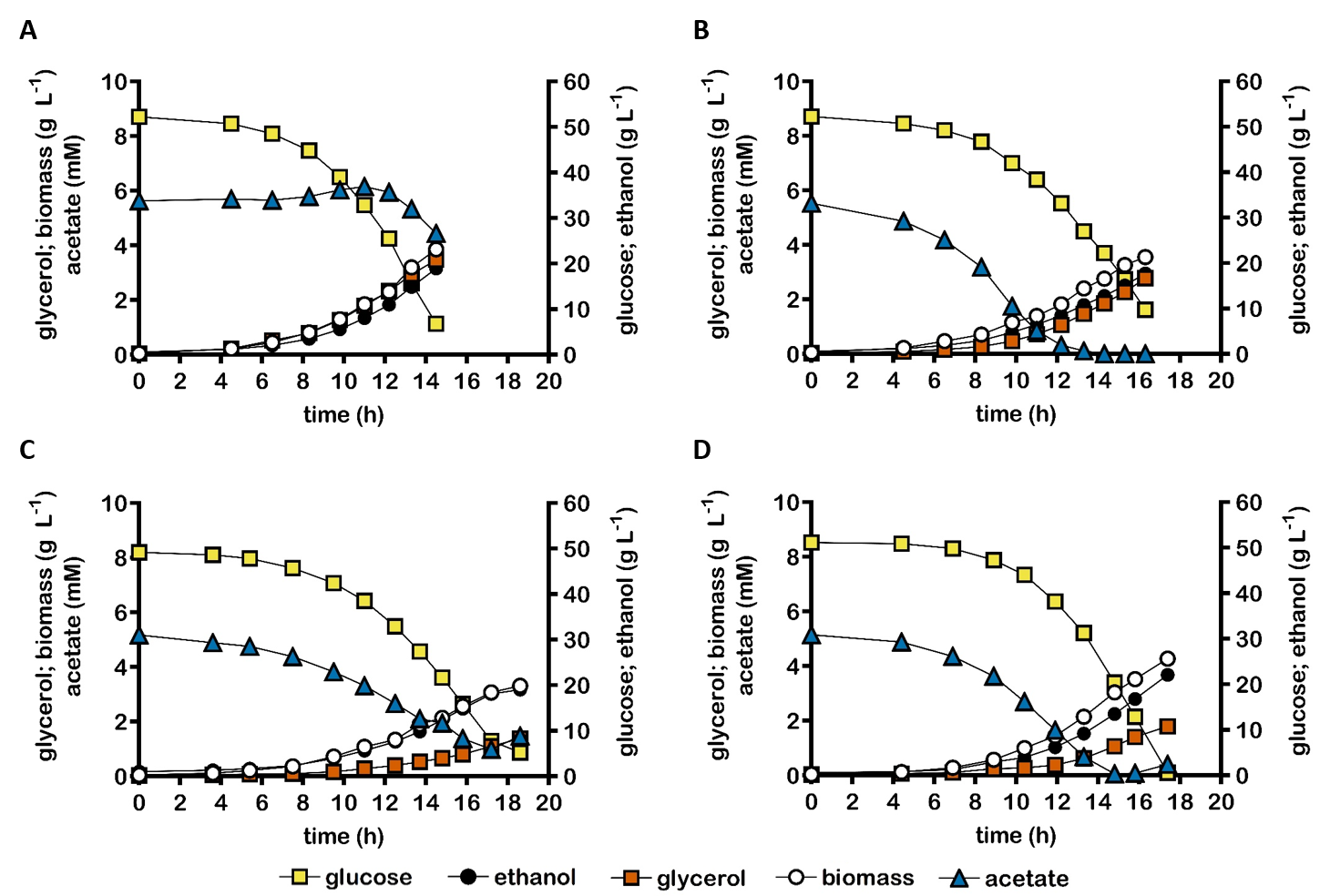

**Figure S3:** Concentrations of biomass, ethanol, glycerol and acetate in anaerobic bioreactor batch cultures of S. cerevisiae strains IME324 (reference strain, **A**), IMX2503 (Δgpd2 Δald6 eutE, **B**), IMX2502 (non-ox PPP↑ Δgpd2 Δald6 pTDH3-eutE pDAN1-prk 2x pTDH3-cbbm pTPI1-groES pTEF1-groEL, **C**), and IMX2723 (non-ox PPP↑ Δgpd2 Δald6 pTDH3-eutE pDAN1-prk-19aa (4) 2x pTDH3-cbbm pTPI1-groES pTEF1-groEL, **D**). Cultures were grown anaerobically at pH 5.0 and at 30 °C on synthetic medium containing 50 g L^-1^ glucose and 5.0 mmol L^-1^ acetate. Non-ox PPP↑ indicates the integration of the expression cassettes of pTDH3-RPE1, pPGK1-TKL1, pTEF1-TAL1, pPGI1-NQM1, pTPI1-RKI1 and pPYK1-TKL2. Representative cultures of independent duplicate experiments are shown, corresponding replicate of each culture shown in Figure 2.
